# Chiral cilia orientation in the left-right organizer

**DOI:** 10.1101/252502

**Authors:** Rita R. Ferreira, Guillaume Pakula, Lhéanna Klaeyle, Hajime Fukui, Andrej Vilfan, Willy Supatto, Julien Vermot

## Abstract

Chirality is a property of asymmetry between an object and its mirror image. Most biomolecules and cells are intrinsically chiral. Whether cellular chirality can be transferred to asymmetry at the tissue scale remains an unresolved issue. This question is particularly relevant in the left-right organizer (LRO), where cilia motility and chiral flow are thought to be the main drivers of left-right axis symmetry breaking. Here, we built a quantitative approach based on live imaging to set apart the contributions of various pathways to the spatial orientation of cilia in the Kupffer’s vesicle (KV, zebrafish LRO). We found that cilia populating the zebrafish LRO display an asymmetric orientation between the right and left side of the LRO. Cilia orientations, therefore, give the KV cells a sense of chirality which is different from the chirality of cilia rotation. Surprisingly, we found this asymmetry does not depend on the left-right signalling pathway or flow. Furthermore, we show the establishment of the chirality is dynamic and depends on planar cell polarity. Together, this work identifies a different type of asymmetry in the LRO and sheds light on the complexity of chirality genesis in developing tissues.

---

A chiral object cannot be superimposed on its mirror image. Most biological molecules are chiral. Yet, how macroscopic chiral asymmetries arise in physics and biology is still debated (Morrow et al., 2017; Wagnière, 2007). In living systems, a number of independent mechanisms of chirality establishment have been identified, from the subcellular to the tissue scale (Blum et al., 2014; Coutelis et al., 2014; Dasgupta and Amack, 2016; Gomez-Lopez et al., 2014; Hamada and Tam, 2014; Levin, 2005; Naganathan et al., 2014; Noel et al., 2013; Tee et al., 2015). The most studied system is certainly the mechanism that sets asymmetric gene expression around the left-right organizers (LRO) of vertebrates (Ferreira and Vermot, 2016. Generally, asymmetrical signals are generated in LROs as a response to a directional flow driven by motile cilia (Shinohara and Hamada, 2017). In the LRO, the orientation of cilia rotation follows a clockwise motion and is invariable amongst vertebrates (Okada et al., 2005). These properties are key for controlling the chiral direction of the flow (Hilfinger and Julicher, 2008; Shinohara and Hamada, 2017). Nevertheless, other cellular chiral features at the scale of the whole LRO have never been investigated.

In zebrafish, the LRO is called the Kupffer’s vesicle (KV) (Figure 1A). Before any sign of asymmetric cell response, the KV consists of a sphere containing monociliated cells where a directional flow progressively emerges as a result of stereotyped cilia spatial orientation (Ferreira et al., 2017). Over the course of hours, the cilia-generated flow triggers an asymmetric calcium response on the left side of the cavity (Francescatto et al., 2010; Sarmah et al., 2005; Yuan et al., 2015), and, consequently, a left-biased asymmetric pattern of gene expression (Essner et al., 2005; Kramer-Zucker et al., 2005). Coordinating appropriate cilia spatial orientation with directional flow generation is thus critical for the subsequent asymmetric response and proper LR patterning (Hashimoto and Hamada, 2010). Current studies have posited that flow patterns arise first in the LRO and then dictate the symmetry-breaking event (Blum et al., 2014; Shinohara and Hamada, 2017). This has led to the inference that symmetry breaking initiation depends on the establishment of symmetrical LRO where cilia orientation is tightly controlled by the Planar Cell Polarity (PCP) pathway (Hashimoto and Hamada, 2010; Marshall and Kintner, 2008; Song et al., 2010). In this model, the direction of cilia rotation leading to the directional flow is the only known chiral element in the LRO. However, a number of studies have suggested that subcellular chirality associated with cytoskeletal asymmetric order could also participate in setting the LR axis, in particular in asymmetric animals where no LRO have been identified (Davison et al., 2016; Hozumi et al., 2006; Kuroda et al., 2009; Sato et al., 2015; Shibazaki et al., 2004; Speder et al., 2006). This raises the intriguing possibility that the LRO could use subcellular chiral information for symmetry breaking. In the absence of tools for visualizing potential chirality in the LRO, however, it is difficult to establish if cell chirality could participate in the process of symmetry breaking. To meet this challenge, we developed a quantitative analysis based on live imaging allowing the investigation of the LRO chirality and the identification of the factors controlling it.

**Figure 1:**
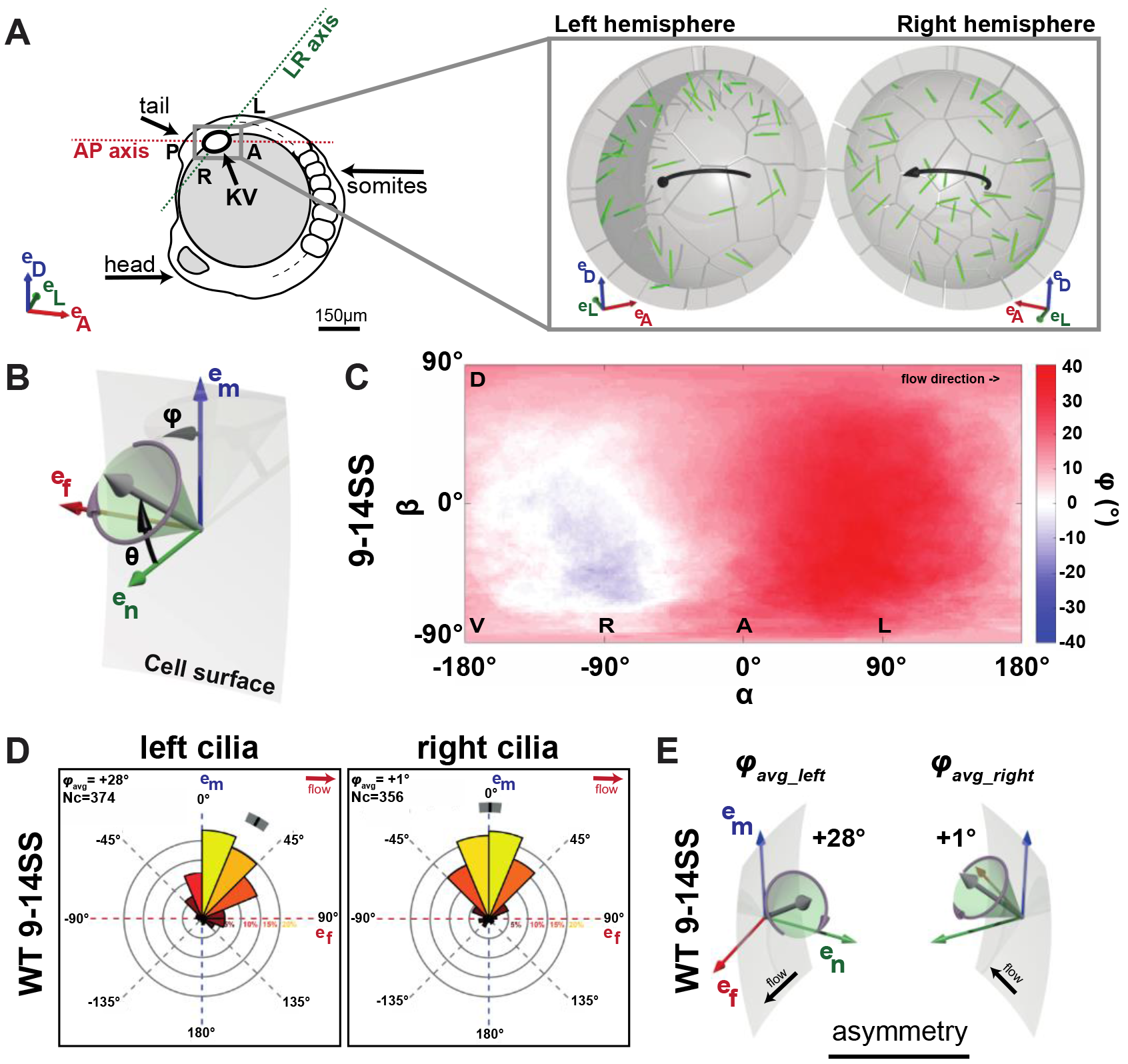
The zebrafish left-right organizer is asymmetric. **(A)** Schematics of the zebrafish embryo (left panel) highlighting the KV localization (grey box). The zoom-up box (right panel) shows the transverse section of the KV, depicting the cilia (in green) and the directional flow (black arrows). The embryonic body plan axes are marked as AP (anterior-posterior) and LR (left-right). The body plan reference frame is defined by basis vectors e_D_ (dorsal), e_L_ (left), e_A_ (anterior). **(B)** Cilia orientations are represented by two angles: θ (tilt angle from the surface normal e_n_) and φ (angle between the surface projection of the ciliary vector and the meridional direction e_m_). Cell surface is represented in grey, e_m_ in blue, e_f_ in red and the normal e_n_ in green. **(C-D)** Distributions of φ at 9-14SS in a 2D flat map (C) or rosette plots (D) obtained from 14 wild-type vesicles with a total of 730 cilia. (C) Average 9 values displayed in a 2D flat map showing 9 on the left (0° ≤ α ≥ 90° and −90° ≤ β ≥ 90°) is higher (red) than φ on the right (−90° ≤ α ≥ 0° and −90° ≤ β ≥ 90°). **(D)** Rosette plots showing the φ angle distribution for the left and right-sided motile cilia, and the 95% confidence interval (grey stripe) for the population mean (black tick). In each rosette, 0° indicates the meridional direction (e_m_) and 90° the flow direction (e_f_). Most cilia exhibit φ angles between [−45°; +45°], corresponding to a meridional tilt (Ferreira et al., 2017). **(E)** Schematics showing the φ_avg_ of the 3D resultant vector on the left and right sides of the KV at 9-14SS. Nc = number of cilia.

Tissue chirality can result from asymmetric cell shape and asymmetric organelle distribution at the cell scale (Wan et al., 2011; Xu et al., 2007). We reasoned that as an asymmetry generator, the LRO itself constitutes a candidate for being a chiral organ. We made use of the ellipsoidicity of the KV to assess the symmetry of cilia orientation by focusing on the two angles defining cilia orientation in 3D (Figure 1B): θ (tilt) is the angle of the cilium with respect to the KV surface normal (0° for a cilium orthogonal to the KV surface and 90° for parallel); φ is the orientation of the cilium projected on the KV surface (0° for a cilium pointing in a meridional direction towards the dorsal pole). Thus, the meridional tilt of cilia reported in wild-type (WT) KV corresponds to θ > 0° and φ close to 0°, meaning that cilia point dorsally following the meridians of a sphere (Ferreira et al., 2017). We performed live-imaging using the zebrafish *act2b:Mmu.Arl13bGFP* transgenic line where cilia are fluorescently labelled (Borovina et al., 2010). Next, we extracted the angles θ_avg_ and φ_avg_ of the average cilia orientation vector in both hemispheres of the KV between the 9 and 14 somite stage (SS), when the chiral flow is fully established. To quantify differences in cilia orientation between the left and right sides, we calculated separately for each side the average cilium direction in local coordinates (Figure 1B). In analogy to an individual cilium, the angles φ_avg_ and θ_avg_ describe the direction of the average orientation vector on each side. In case of a mirror-symmetric KV, θ_avg_ of the average cilium is equal on the left and on the right sides and the φ_avg_ angles are mirror-imaged: θ_avg_left_ = θ_avg_right_, and φ_avg-left_ = − φ_avg_right_ = φ_avg_right_mirror_. To quantitatively assess the significance of asymmetries in the KV, we designed a permutation test based on the definition of chirality (see Methods) and calculated p-values estimating the likelihood that an *a priori* symmetric KV will show an equal or larger difference |φ_avg_left_ − φ_avg_right mirror_| due to variability.

We extracted and averaged the results obtained from 14 WT vesicles with a total of 730 cilia, and estimated the θ_avg_ and φ_avg_ angles of the average cilium in 3D. There was a difference between the measured φ angle on the left and the right sides of the KV (Figure 1C). While right-sided cilia are almost perfectly oriented along the meridional direction (φ_avg_right_ = +1°, Figure 1D-E), cilia in the left hemisphere exhibit a strong tilt following the direction of the flow (φ_avg_left_ = +28°, Figure 1D-E). On average, it defines a dextral orientation over the whole vesicle (Sup. Figure 2A). The permutation test confirmed the significance (p<0.001) of the observed asymmetry in cilia orientations between the left and right side of the KV (φ_avg_left_ ≠ − φ_avg_right_) (Methods and Table 1). No difference in tilt angles was observed between the left and right hemispheres (θ_avg_left_ = θ_avg_right_; Sup. Figure 1A). Together, these results show that cilia orientation is asymmetric in the KV and follows a dextral chirality.

**Table 1:** Statistical test of mirror-symmetry in the KV. The p-values result from a permutation test under the null hypothesis of a mirror-symmetric KV φ_avg_left_ = − φ_avg_right_.

| Mirror-symmetry: $\varphi_{\text{Left}} = -\varphi_{\text{right}}$? | WT 3SS | | WT | WT | WT | *dnaaf1*<sup>-/-</sup> | *dnaaf1*<sup>-/-</sup> | Blebbistatin | *rock2b*-MO | *cup*<sup>-/-</sup> | *spaw*<sup>-/-</sup> | *trilobite*<sup>-/-</sup> |
| --- | --- | --- | --- | --- | --- | --- | --- | --- | --- | --- | --- | --- |
|  | motile cilia | immotile cilia | 6SS | 8SS | 9-14SS | 3SS | 8SS | 8SS | 8SS | 8SS | 8SS | 8SS |
| p value | 0.56 | 0.23 | 0.082 | 0.003 | <0.001 | 0.23 | <0.001 | <0.001 | 0.014 | <0.001 | <0.001 | 0.54 |

To gain a better sense of when the chirality of cilia orientation begins, we analyzed cilia orientation in KV in embryos at different developmental stages. We quantified the φ angle distributions of cilia from both hemispheres of WT embryos at 3SS (early-stage), when the first signs of LR asymmetry have been reported (Yuan et al., 2015), at 6SS and at 8SS (mid-stages). At 3SS, two populations of cilia exist in the KV, motile and immotile. Neither the motile (φ_avg_left_ = +15° and φ_avg_right_ = −20°; Figure 2A-B), nor the immotile cilia population (φ_avg_left_ = +32° and φ_avg_right_ = −8°; Sup. Figure 2B) exhibit a significant difference between the left and right sides of the KV (φ_avg_left_ ≈ − φ_avg_right_, Table 1), resulting in a φ_avg_ close to 0° at 3SS (Figure 3A). WT embryos at 6SS show some asymmetry (Figure 3A), but it is not yet statistically significant (p=0.086, Table 1). Interestingly, at 8SS the side-biased orientation (φ_avg_left_ = +19° and φ_avg_right_ = +4°; Figure 2A-C) becomes significant enough to reveal an overall asymmetry of the KV (Figure 3A and Table 1). These results show that the orientation differences are not changing linearly between left and right, where the left angle does not change much between 3SS and 8SS and the right side changes more significantly. We also assessed the variability of cilia orientation by plotting the mean orientation on the left vs. right side for each embryo (Sup. Figure 3A). The variability between embryos is always substantial, but it reduces with time and all 14 embryos at 9-14SS show asymmetry in the same direction. When considering θ angle distributions at 3, 6 and 8SS, we did not find any asymmetry between the left and right-side of the KV, demonstrating that the cilia tilt remains symmetrical over time (Sup. Figure 1A). Together, these results demonstrate that cilia orientation in the LRO does not exhibit any asymmetry until 6SS and becomes progressively asymmetric during the course of KV development and LR patterning.

**Figure 2:**
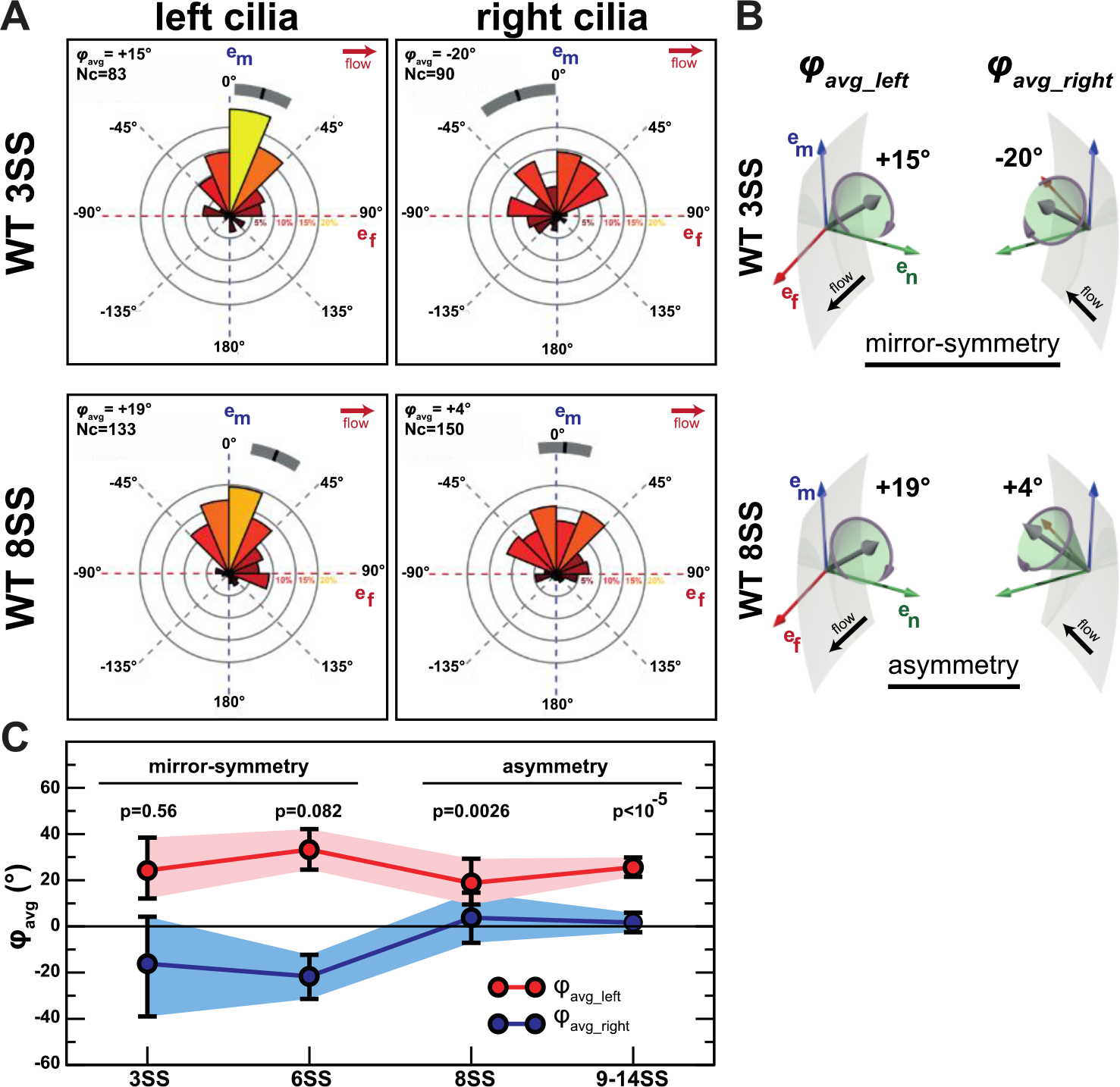
Asymmetric cilia orientation arises over time. **(A)** Rosette plots showing the φ angle distributions for the left- (left panel) and right-sided (right panel) cilia, with their mean (black tick) and associated 95% confidence interval (grey stripe), for WT 3SS (upper panel) and WT 8SS (lower panel). In each rosette, 0° indicates the meridional direction (e_m_) and 90° the flow direction (e_f_). **(B)** Schematics showing the φ_avg_ of the 3D resultant vector on the left and right sides of the KV at 3SS (upper panel) and 8SS (lower panel). **(C)** Mean orientation φ_avg_ of motile cilia in the left (red) and right (blue) half of the KV as a function of time. The error bars and shaded regions display 95% confidence intervals. Nc = number of cilia.

**Figure 3:**
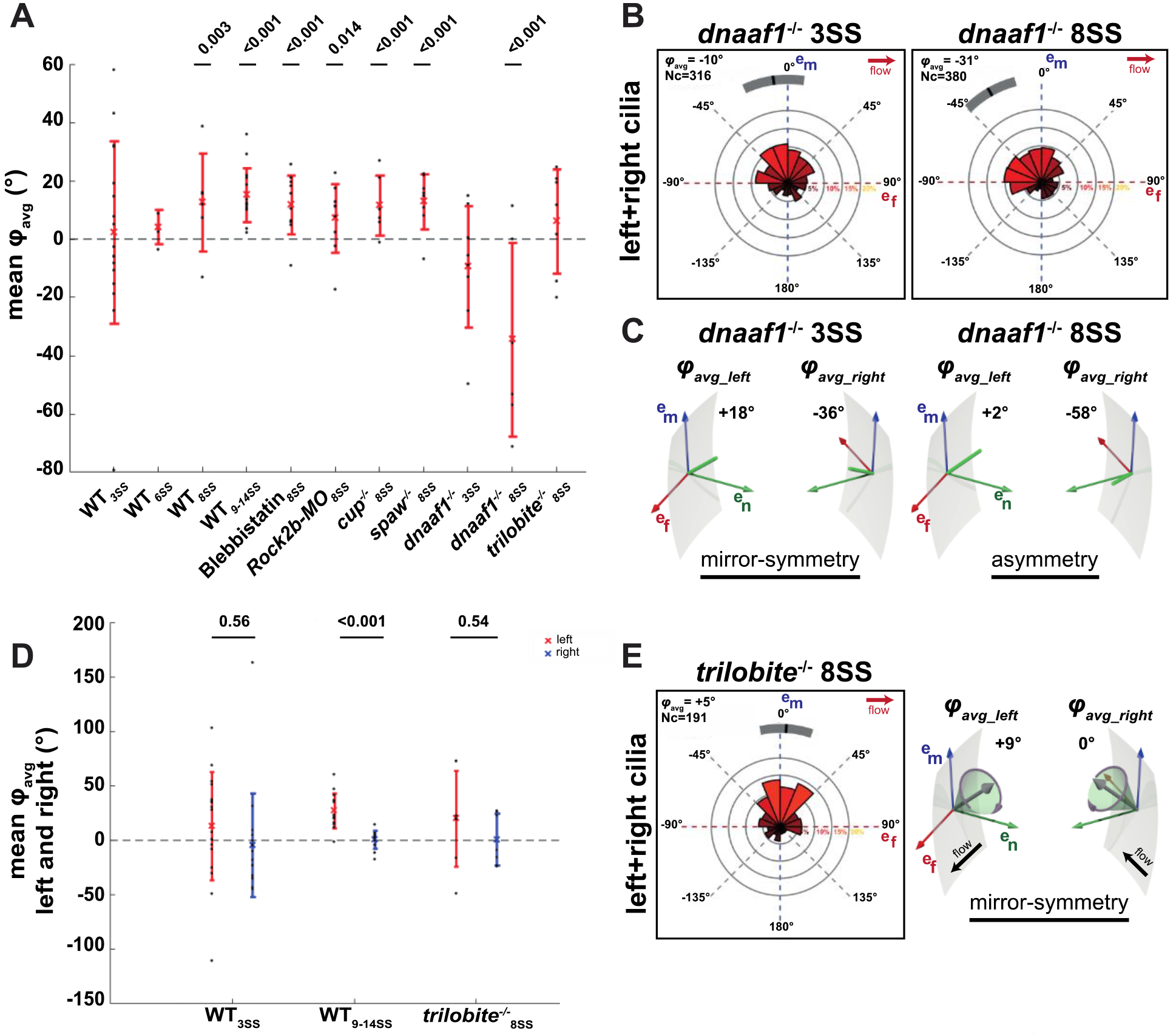
DNAAF1 and PCP are important for asymmetric cilia orientation. **(A)** Dot plot displaying the mean values of φ_avg_ for each condition. Each black dot represents the mean values for one individual KV, the cross displays the average value for all KVs of the respective condition and the line the standard deviation. The mean values of φ_avg_ are always greater than zero, except for *dnaaf1*^−/−^, which are below zero, revealing that its orientation with respect to the meridional direction is reversed. Also, WT_3SS_, WT_6SS_, *dnaaf1*_3SS_ and *trilobite*^−/−^_8SS_ have φ_avg_ mean values close to zero, demonstrating a LR symmetry in the KV. The p-values result from a permutation test under the null hypothesis of a mirror-symmetric KV φ_avg_left_ = − φ_avg_right_, which is equivalent to φ_avg_ = 0 (see Methods). **(B)** Rosette plots showing the φ angle distribution of cilia in both KV hemispheres (left+right cilia), for *dnaaf1*^−/−^ 3SS (upper panel) and *dnaaf1*^−/−^_8SS_ (lower panel). Both *dnaaf1*^−/−^ 3SS and *dnaaf1*^−/−^ 8SS cilia are inclined in the opposite direction to the flow (e_f_). **(C)** Schematics showing the φ_avg_ of the 3D resultant vector on the left and right sides of the KV for *dnaaf1*^−/−^ 3SS (upper panel) and *dnaaf1*^−/−^ 8SS (lower panel). **(D)** Dot plot displaying the mean values of φ_avg_ on the left (in red) and on the right (in blue) sides of the KV for WT_3SS_, WT_9-14SS_, and *trilobite*^−/−^_**8SS**_. *trilobite*^−/−^_8SS_ display no difference between the mean φ_avg_ on the left and right sides of the KV. The p-values result from a permutation test under the null hypothesis of φ_avg_left_ = − φ_avg_right_. **(E)** Rosette plot showing the φ angle distribution of cilia in both KV hemispheres (left+right cilia) for *trilobite*^−/−^ 8SS. In contrast to other conditions studied so far, φ_avg_ of *trilobite*^−/−^ KV cilia is close to zero, reinforcing the observation that the KV is symmetrical at 8SS. **(F)** Schematics showing the φ_avg_ on the left and right sides of the KV for *trilobite*^−/−^_8SS_. Nc = number of cilia.

Considering the function of the KV is to generate an asymmetrical signal, we investigated whether the observed asymmetry of the KV itself is modulated by the LR signalling pathway. According to current models of symmetry breaking, the asymmetric expression of genes around the LRO is under the control of *pkd2* that acts as a flow sensor and leads to the left-sided expression of *spaw*, a nodal-related gene involved in the establishment of the left-right asymmetry (Schottenfeld et al., 2007; Yuan et al., 2015). To test the impact of flow sensing and asymmetric gene expression downstream of flow on asymmetric cilia orientation, we used *cup*^*tc241*^ (Schottenfeld et al., 2007) and *spaw*^*s457*^ (Kalogirou et al., 2014) mutants where *pkd2* and *spaw* are not functional, respectively. As expected, both *cup*^−/−^ and *spaw*^−/−^ embryos have laterality defects (Sup. Figure 4B-C and Table 2). By analyzing the KV cilia orientation, we found a normal meridional orientation of cilia (Sup. Figure 2C, φ_avg_ ≠ 0° in Figure 3A) and permutation tests show that there is an overall asymmetry in the KV (φ_avg_left_ ≠ − φ_avg_right_) in both *cup*^−/−^ and *spaw*^−/−^ KV cilia at 8SS. Together these results indicate that asymmetric cilia orientation is not dependent upon the LR signalling cascade in the LRO.

**Table 2:** Relative frequencies of LR scoring outcomes and their standard errors (Supplemental Figure 4).

|  | WT controls | blebbistatin | rock2b MO | *dnaaf1*<sup>-/-</sup> | *cup*<sup>-/-</sup> | *spaw*<sup>-/-</sup> | *trilobite*<sup>-/-</sup> |
| --- | --- | --- | --- | --- | --- | --- | --- |
|  | Mean ± SEM | Mean ± SEM | Mean ± SEM | Mean ± SEM | Mean ± SEM | Mean ± SEM | Mean ± SEM |
| left | 92.3 ± 4.0% | 71.2 ± 2.5% | 27.9 ± 1.3% | 16.5 ± 5.1% | 0 ± 0% | 0 ± 0% | 46.5 ± 7.3% |
| bilateral | 7.7 ± 4.0% | 17.1 ± 4.8% | 48.1 ± 4.0% | 65.2 ± 11.1% | 98.8 ± 1.2% | 0 ± 0% | 53.2 ± 6.9% |
| right | 0 ± 0% | 4.2 ± 1.1% | 6.9 ± 0.5% | 16.1 ± 5.1% | 1.2 ± 1.2% | 0 ± 0% | 0 ± 0% |
| absent | 0 ± 0% | 7.5 ± 1.3% | 17.1 ± 5.9% | 2.2 ± 1.3% | 0 ± 0% | 100 ± 0% | 0.3 ± 0.4% |
| *situs solitus* | 93.4 ± 0.6% | 68.3 ± 4.3% | 46.3 ± 3.0% | 35.1 ± 1.4% | 25.5 ± 4.6% | 16.5 ± 0.6% | 23.5 ± 0.2% |
| *heterotaxy* | 5.3 ± 0.9% | 28.3 ± 2.9% | 40.9 ± 1.8% | 39.6 ± 0.6% | 63.5 ± 4.6% | 76.3 ± 1.7% | 63.5 ± 0.6% |
| *situs inversus* | 1.3 ± 0.3% | 3.5 ± 1.3% | 12.9 ± 1.2% | 25.3 ± 1.2% | 11.5 ± 0.4% | 7.2 ± 2.3% | 13.0 ± 0.4% |

**Table 3:**
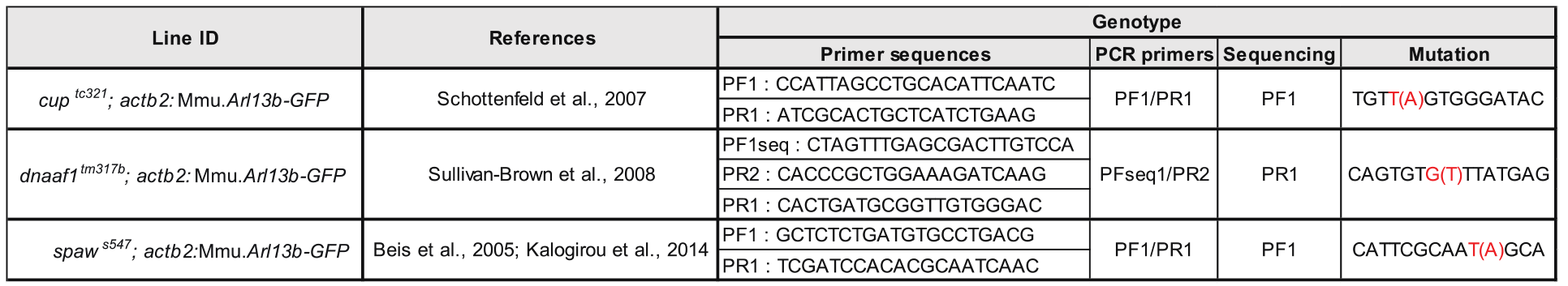
Supplemental information concerning the genotyping strategies of the mutant lines used, including the designed primers and mutation details. All primers were designed with the program ApE (http://biologylabs.utah.edu/jorgensen/wayned/ape/) and using the genomic sequences available on Ensemble (*Ensembl genome browser 84*) for *dnaaf1, spaw* and *cup* genes. Embryos from the *trilobite*^*tc240a*^ (Heisenberg and Nusslein-Volhard, 1997) mutant line were identifiable by phenotype at all the stages analyzed. *dnaaf1*^*tm317b*^ (Sullivan-Brown et al., 2008) *and cup*^*tc241*^ (Schottenfeld et al., 2007) mutant embryos were identifiable for heart and gut scoring analysis (48-53 hpf) and were genotyped only at earlier stages. *spaw*^*s457*^ (Kalogirou et al., 2014) mutant embryos were always genotyped by sequencing.

Actomyosin contractility is a well-known modulator of tissue chirality *in vitro* (Wan et al., 2011) and *in vivo* during early development of asymmetry in Xenopus embryos (Qiu et al., 2005), and cardiac looping in vertebrate embryos (Noel et al., 2013; Taber, 2006). In addition, it has been shown that the migration of the basal body to the apical surface of cells is essential for cilia formation (Hong et al., 2015; Pitaval et al., 2010) as well as important for cell-cell tension regulation during KV morphogenesis (Wang et al., 2012) and LR determination (Gros et al., 2009; Tabin and Vogan, 2003; Wang et al., 2012). Furthermore, actomyosin contractility has been shown to be important for KV morphogenesis and flow (Wang et al., 2011; Wang et al., 2012). We thus assessed its impact on asymmetric cilia orientation. To do so, we used blebbistatin and the *rock2b*-morpholino (*rock2b*-MO) to block myosin-II activation (Wang et al., 2011). Both blebbistatin-treated and *rock2b*-MO embryos exhibit laterality defects (Sup. Figure 4B-C and Table 2) and abnormal cell clustering in the anterior side of the KV (Sup. Figure 1D) as previously described (Wang et al., 2011; Wang et al., 2012). Cilia orientation analysis showed that the *rock2b*-Myosin-II pathway does not interfere with the meridional tilt of KV cilia (Sup. Figure 2C). Permutation tests show that the KV at 8SS is overall asymmetric (φ_avg_left_ ≠ − φ_avg_right_) for both blebbistatin and *rock2b*-MO treated embryos (φ_avg_ ≠ 0° in Figure 3A with corresponding p values; Table 1). Overall these results indicate that asymmetric cilia orientation is not under the control of the *rock2b*-Myosin-II pathway (Ferreira et al., 2017), and, more generally, the main components of the LR signaling acting upstream and downstream of the flow operating in the LRO.

Ciliary components involved in cilia motility have also been shown to modulate cilia orientation (Jaffe et al., 2016). We thus assessed if cilia motility could be a factor controlling asymmetric cilia orientation. We first analyzed cilia orientation in the WT at 3SS and found that WT immotile cilia have a distinct orientation compared to the motile cilia at the same stage (θ_avg_WT3SS immotile_ = +15° and θ_avg_WT3SS motile_ = +30°; p<10^−4^), suggesting that motility could be involved in modulating cilia orientation. To confirm cilia motility involvement, we analyzed the effects *dnah9* (*lrdr1*) knock-down and found that asymmetric cilia orientation was perturbed (FigureS2A). We next studied the *dnaaf1*^−/−^ *(dynein axonemal assembly factor 1;* old nomenclature: *lrrc50* - *leucine-rich repeat-containing protein 50)* mutants. *dnaaf1* encodes for a cilium-specific protein required for the stability of the ciliary architecture and when mutated abrogates its ability to interact with specific targets important for cilia motility (Sullivan-Brown et al., 2008). Importantly, *dnaaf1*^−/−^ cilia have ultrastructural defects and display abnormal dynein arms orientation in beating cilia (Loges et al., 2009). We found that all cilia are immotile in the *dnaaf1^−/−^* KVs (Movie 1) and that LR axis establishment is randomized (Sup. Figure 4B-C and Table 2). We next assessed the symmetry in the *dnaaf1^−/−^* cilia and found that their orientation does not show any sign of asymmetry at 3SS (φ_avg_left_ ≈ −φ_avg_right_) (φ_avg_ = 0° in Figure 3A; Figure 3B-C and Table 1), similar to the controls at the same stage (Figure 2 and Sup. Figure 2B). At 8SS, we found that *dnaaf1^−/−^* cilia orientation is asymmetric (φ_avg_left_ ≠ −φ_avg_right_ in Figure 3B-C and Table 1; φ_avg_ ≠ 0° in Figure 3A) showing that asymmetric cilia orientation is flow independent. Surprisingly though, we found that *dnaaf1^−/−^* cilia have a meridional tilt (Figure 3B) but are inclined in a sinistral direction, which is the opposite direction than WT at 8SS (φ _avg_dnaaf1_^−/−^8SS = −31° (Figure 3B) and φ_avg_WT8SS motile_ = +11° (Sup. Figure 2A); p<10^−5^. See also Figure 3A with φ_avg_ < 0°). Thus *dnaaf1* is involved in the control of chiral cilia orientation and it may do so independently of the flow.

Finally, since cilia motility and planar cell polarity are interdependent (Jaffe et al., 2016), we directly tested the role of the PCP in KV chirality. It has been shown that the PCP pathway modulates cilia tilt in the mouse LRO (Hashimoto and Hamada, 2010; Marshall and Kintner, 2008; Song et al., 2010). Generically, PCP proteins are required to establish cell polarity within tissues across a large variety of animal species (Wallingford, 2012). We used the zebrafish *trilobite* mutant line, where Van Gogh/Strabismus homologue - *Van gogh-like 2* (*Vangl2*) -, a gene encoding a protein essential for PCP signalling, is mutated (Jessen and Solnica-Krezel, 2004). *Vangl2* function is required for the posterior tilt observed in KV cilia, and anomalies in cilia orientation disrupt the cilia-driven flow and LR determination (Borovina et al., 2010). After confirming the *trilobite*^*tc240a*^ have left-right defects (Sup. Figure 4B-C and Table 2), we studied cilia orientation at 8SS using *trilobite*^*tc240a*^; *actb2:Mmu.Arl13b-GFP* (Heisenberg and Nusslein-Volhard, 1997) mutant embryos, and found cilia orientation still follows a meridional tilt (Figure 3E and Sup. Figure 1A). Our flow simulations (see Methods and Sup. Figure 4D) predict that flow is significantly weaker in *trilobite*^*tc240a*^ than in the WT (p=0.024), as expected for a PCP mutant (Borovina et al., 2010). In contrast with other conditions studied so far, permutation tests could not detect any asymmetry in the *trilobite*^−/−^ KV cilia at 8SS, meaning φ_avg_left_=−φ_avg_right_ (p=0.54, Figure 3D,E, Sup. Figure 2C and Table 2) with a φ_avg_ ≈ 0° (Figure 3A). Altogether, these results indicate that the PCP pathway is involved in chiral cilia orientation in the KV.

Since PCP is known to alter the basal body position at the cell surface of the vertebrate LRO (Borovina et al., 2010; Hashimoto and Hamada, 2010; Juan et al., 2018; Song et al., 2010), we developed a quantitative analysis of basal cilia orientations in the KV to test if both are interdependent (Sup. Figure 3B-C). As expected, we found that the basal body is localized posteriorly along the anterior-posterior (AP) axis of the WT embryo and is symmetrical along the left-right axis of the cell (Figure 4A, C, D). We next assessed the AP position of the cilia in the *dnaaf1* mutants and found that it is altered along the AP axis, with basal body located more centrally to the cell (Figure 4 A-D), in line with the finding that Dnaaf1/Lrrc50 interacts physically with C21orf59, which is involved in polarizing motile cilia (Jaffe et al., 2016. By contrast, we did not detect any chirality in the basal body AP or right-left (RL) positions when testing the positions of right cilia against the mirror image of left cilia in controls and mutants (p>0.15). Thus, the chirality of cilia orientation seems not dependent on basal body positioning along the LR axis of the cells, which more generally suggests cilia orientation is not solely controlled by basal body position. In addition, these results show that *dnaaf1* is involved in setting AP basal position and that cells might need to be properly planar polarized for the establishment of chiral cilia orientation. Overall, we conclude that proper cilia positioning along the AP axis of the cell and asymmetric cilia orientation might be interdependent.

**Figure 4:**
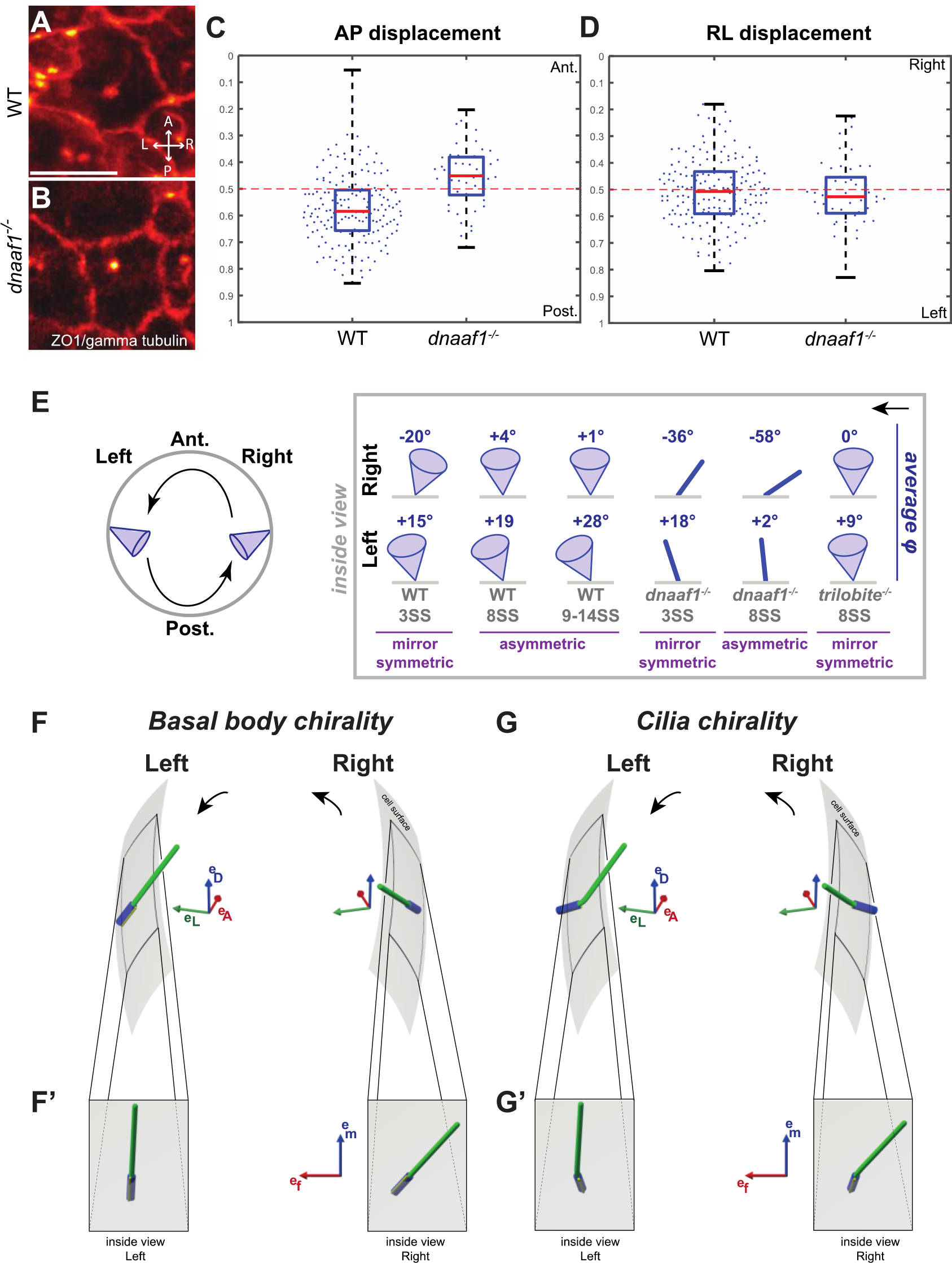
Basal body position in KV cells, schematic summary of the contributions of different pathways and the potential origins of chirality leading to asymmetric cilia orientation in the KV. **(A-D)** Basal body position analysis. The positions of the cilia basal body relative to the AP (Anterior-to-posterior) or LR (Left-to-right) extension of each cell were extracted from WT **(A)** or *dnaaf1*^−/−^ embryos **(B)** using an orthogonal projection of the fluorescence intensity on the plane tangential to the KV surface after ZO1 and gamma tubulin staining of KV cells. The distributions of AP displacement (dots from 0 for anterior to 1 for posterior in **(C)**) and LR displacement (dots from 0 for right to 1 for left in **(D)**) are centered at the middle of the cell (displacement at 0.5) for all cases, except a significant posterior displacement of basal bodies in the WT. **(E)** Left: Dorsal view of a mirror-symmetric KV showing a representative cilium on the left and on the right side. Right: schematic view of the average φ orientation over time in WT, *dnaaf1*^−/−^ and *trilobite*^−/−^ KV for cilia in the right (upper row) and in the left (lower row) KV hemispheres. KV is mirror-symmetric at 3SS both in WT and *dnaaf1*^−/−^ embryos evolving to an asymmetric orientation at 8SS. In contrast, *trilobite*^−/−^ 8SS KVs do not show any evidence of asymmetry. **(F-G)** Schematics of the two hypotheses for the observed chirality (dorsal-posterior views of the KV in F and G, inside views of both sides in F’ and G’): **(F)** the orientation of cilia can be asymmetric between left and right because the orientation of the basal bodies is chiral; **(G)** alternatively, the basal bodies can be arranged symmetrically, but the intrinsic chiral structure of each cilium leads to an overall asymmetric distribution of cilia orientations. The black arrows in E, F and G indicate the direction of the flow.

## Discussion

Previous work has focused on identifying signals that trigger the specification of the left embryonic side in response to flow, but the chirality of the LRO remained untested. We took advantage of the zebrafish LRO (KV) as an established model system in which cilia orientation and LR symmetry can be accurately quantified and show that cilia orientation is not symmetrical between the left and right side of the LRO. Our study revealed that cilia orientation progressively changes from mirror-symmetric to asymmetric (Figure 4E) and that this new type of asymmetry emerges independently of the LR symmetry cascade. Interestingly, even if we found that there is a strong variability in cilia orientation, we found that they will invariably become chiral in every KV, suggesting a very robust process for chirality determination.

Considering that several key modulators of symmetry breaking such as Pkd2, Rock2b (and the associated actomyosin pathway) or Spaw do not affect cilia orientation, we conclude that the LRO promotes left-right symmetry breaking and asymmetric cilia orientation through distinct mechanisms. If the asymmetry of cilia orientation is independent of the LR machinery, how can we explain its emergence? One possibility would be that cilia reorient themselves in response to the flow so that the flow forces themselves would be the source of chiral information. During blood vessel morphogenesis, endothelial cells polarize themselves against the flow in a similar way (Franco et al., 2015; Kwon et al., 2016. There are, however, three strong arguments opposing this hypothesis. First, we still observe an asymmetric orientation of cilia in *dnaaf1* mutants where flow is absent and in *rock2b* knockdown where flow is weak. Second, our previous work (Ferreira et al., 2017) along with our simulations (Sup. Figure 4E) show that the torque resulting from the global flow is much smaller than the drag on a motile cilium. A similar conclusion has been drawn when comparing the forces exerted on cells by beating ependymal cilia with those mediated by the fluid (Mahuzier et al., 2018). Finally, we found that the left cilia reorient less than the right cilia even though the flow has the same magnitude on both sides (Ferreira et al., 2017. It seems thus that the relationship between flow direction and cilia orientations cannot be causal. It is more likely that both could reflect disorders in cilia orientation in the respective mutants.

Rather than being induced by the flow, we propose that the asymmetry could arise from chiral influences generated by the cytoskeletal components that operate at the cellular and subcellular scales (Satir, 2016). In line with this idea is the fact that cells can display chiral behaviours *in vitro* and *in vivo* independently of a LRO (Naganathan et al., 2014; Noel et al., 2013; Speder et al., 2006; Tee et al., 2015). For example, basal body and cilia ultra-structure display obvious signs of chirality (Afzelius, 1976; Marshall, 2012; Pearson, 2014). This suggests two possibilities: either the orientation of the cilia basal bodies is chiral (asymmetric between left and right, Figure 4F-4F’), or they are oriented symmetrically and the intrinsic chiral structure of each cilium leads to an overall asymmetric distribution of cilia orientations (Figure 4G-4G’). The fact that *dnaaf1*, a cilia specific protein, can reverse cilia orientation from dextral to sinistral seems to argue for the latter. However, we found that *dnaaf1* is also involved in modulating planar cell polarity, so its function within the cilium remains difficult to assess. Interestingly, we did not detect chirality in the distribution of basal body positions. This suggests that the chirality is related to the basal body orientation but not its position. An attractive hypothesis is that cells need to be planar cell polarized to express chirality and that the PCP (through Vangl2 and Dnaaf1) participates in the process that sets basal body orientation. Among the many PCP components that affect LR determination (see for example ciliary components (Jaffe et al., 2016) or unconventional myosin 1d (Juan et al., 2018; Tingler et al., 2018), it will be interesting to assess if some are more important than others in controlling cilia chirality and the robustness of cilia orientation. More work will be needed to establish the molecular basis of cell and cilia chirality in the KV and whether it is conserved in other ciliated LRO.

The demonstration that the asymmetry is actively modulated by the PCP, cilia motility and, potentially, by the internal organization of cilia has important implications for the understanding of chiral information distribution and its control in developing organs. First, it shows that chiral information is dynamic and temporally controlled during the course of LR specification. In that respect, it is interesting that cells establish chiral organization as a result of cell migration *in vitro* (Wan et al., 2011) and that asymmetric cell migration is well described in the chicken LRO (Gros et al., 2009). It is thus possible that the progressive establishment of asymmetric cilia orientation in the LRO reflects an active acquisition of cellular chirality during the time of LRO function. If confirmed, chirality would represent a unique conserved feature between fish and chicken LRO. This raises the intriguing possibility of a more general role of chirality in morphogenetic pattern formation and LR symmetry breaking. Could the chiral cilia orientation participate in symmetry breaking in the LRO? We have previously tested different hypothetical mechanisms of flow detection and shown that a quantitative analysis of physical limits favours chemical sensing (Ferreira et al., 2017). While the basic mechanism of flow generation, as well as flow detection, do not require any prior asymmetry in the KV, we cannot exclude that the ciliary asymmetry participates in the process of symmetry breaking by playing hand-in-hand with the chiral flow to optimize the flow direction or the detection of signalling particles.

More generally, proper cilia orientation is essential for directed flow generation in ciliated tissues or for swimming in ciliated microorganisms (Goldstein, 2015). An important open question is whether cilia can reorient themselves in the direction of flow (Guirao et al., 2010; Mitchell, 2003) and if so, whether this reorientation is a consequence of hydrodynamic forces. Another possibility is that cell polarity is affected by the flow. Since some ciliary proteins involved in cilia motility also participate in planar cell polarity (Jaffe et al., 2016), it is possible that the spatial orientation of motile cilia is an intrinsic mechanism, which is independent of the flow they generate. In the case of brain cavities, the orientation of cilia beating dynamically follows the circadian rhythm and may be driven by transient changes in cell-cell interactions and in PCP (Faubel et al., 2016). Similarly, ciliated microorganisms can reorient the direction of ciliary beating in the course of an avoidance reaction (Tamm et al., 1975). Mechanical strain has also been shown to be involved in dictating cilia orientation, length and motility features (Chien et al., 2018). Our study sheds a different light on these systems as it shows that cilia orientation is related to cell polarity in a complex way that includes an intrinsic sense of chirality. Furthermore, as there is increasing evidence that congenital diseases like idiopathic scoliosis (Grimes et al., 2016), *Kartagener* syndrome, neonatal respiratory distress, hydrocephaly, and male infertility involve cilia motility (Mitchison and Valente, 2017), precise cilia orientation analysis becomes critical to understand the biological principles that govern cilia function and their potential involvement in pathology. In this context, our conclusions and method could be relevant in the studies of a variety of developing organs.

## Materials and Methods

### Zebrafish strains

The zebrafish (*Danio rerio*) lines used in this study were the following: *Tg*(*actb2:Mmu.Arl13b-GFP*) (Borovina et al., 2010), *Tg*(*dnaaf1*^*tm317b*^; *actb2:Mmu.Arl13b-GFP*) (Sullivan-Brown et al., 2008), *Tg*(*trilobite*^*tc240a*^; *actb2:Mmu.Arl13b-GFP*) (Heisenberg and Nusslein-Volhard, 1997), *Tg*(*spaw*^*s457*^; *actb2:Mmu.Arl13b-GFP*) (Kalogirou et al., 2014), *Tg*(*cup*^*tc241*^; *actb2:Arl13b-GFP*) (Schottenfeld et al., 2007). None of the mutant lines display cilia length or KV shape defects (Sup. Figure 1B-C). All zebrafish strains were maintained at the IGBMC fish facility under standard husbandry conditions (14h light/10h dark cycle). Adult fish were anaesthetized with 80μg/mL Tricaine/MS-222 (Sigma, Cat. # A-5040) for genotyping experiments. The Animal Experimentation Committee of the Institutional Review Board of IGBMC approved all animal experiments performed in this project.

### Morpholino (MO) knockdown

MO designed to block the *rock2b* RNA splicing site (Wang et al., 2011) was obtained from *Gene Tools, LLC.* One-cell stage embryos were injected with 0.66ng of *rock2b*-MO (5’-GCACACACTCACTCACCAGCTGCAC-3’).

### Blebbistatin treatment

Embryos were dechorionated and treated with 35μM of Blebbistatin (SIGMA B0560/DMSO) from bud-stage until 3SS when they were washed in 0.3% Danieau medium and kept at 32°C until the desired stage for live imaging (8SS). 1%-DMSO treated embryos were used to monitor potential drug-control effects.

### 2-photon excited fluorescence (2PEF) microscopy

Live imaging experiments were performed as described in (Ferreira et al., 2017), in order to maximize the scanning artefact that allows to properly reconstruct cilia orientation in 3D as described in (Supatto and Vermot, 2011).

### 3D-Cilia Map: quantitative 3D cilia feature mapping

We used *3D-Cilia Map*, a quantitative imaging strategy to visualize and quantify the 3D biophysical features of all endogenous cilia in the Kupffer’s vesicle (KV) in live zebrafish embryos, such as KV size and shape and cilia density, orientation or motility. This image analysis workflow using Imaris (Bitplane Inc.) and custom-made scripts in Matlab (The MathWorks Inc.) was first described in (Ferreira et al., 2017). We improved its automation to facilitate the analysis of a large number of cilia. In addition, we added new feature quantification, such as the length of both motile and immotile cilia, which is estimated based on the radial fluorescence intensity profile originating from the position of each cilium basal body.

All coordinate system definitions are described by (Ferreira et al., 2017). In particular, the cilium orientation is represented as a unit vector from its base to its tip, with angle θ and φ defined in a local basis (Fig 1B). This vector represents the orientation of the rotation axis of motile cilia or of the cilium body orientation in the case of immotile cilia, which are both obtained from experimental images. The average angles φ_avg_ and θ_avg_ used throughout this work describe the direction of the 3D resultant vector, which is the sum of all considered cilia unit vectors.

### Whole-mount *in situ* hybridization (WISH)

Whole-mount *in situ* hybridization was performed as described previously (Thisse and Thisse, 2008). Digoxigenin RNA probes were synthesized from DNA templates of *spaw* (Long, 2003) and *foxA3* (Monteiro et al., 2008). Embryos for *spaw* and *foxA3* WISH were fixed at 17SS and 53 hours post fertilization (hpf) respectively. The zebrafish heart looping was assessed at 48hpf when the heart is already beating. Due to its transparency, the heart loop can be visible using brightfield illumination. We performed WISH for *foxA3* at 53hpf in order to visualize the gut *situs* (Monteiro et al., 2008) in the same embryos in which we previously assessed the heart looping at 48hpf. Embryos were evaluated after WISH and scored according to the curvature between the liver and the pancreas. For the sake of simplicity, we merged the laterality information of both heart and gut and described it according to the clinical terminology: *situs solitus* (heart and gut with normal orientation), *situs inversus* (complete reversal of both organ laterality) and *heterotaxy* (any combination of abnormal LR asymmetries that cannot be strictly classified as *situs inversus*) (Fliegauf et al., 2007; Ramsdell, 2005; Shapiro et al., 2014; Sutherland and Ware, 2009). *Spaw* expression patterns in the lateral plate mesoderm (LPM) can be classified into four main categories: left, bilateral, right or absent (Sup. Figure 4A)(Long, 2003). After scoring, embryos were individually genotyped.

### Immunohistochemistry

Embryos at 8SS were fixed by MEMFA (3.7% formaldehyde, 0.1M MOPS, 2mM EGTA, 1mM MgSO_4_) for 2h at room temperature (RT). Embryos were changed to 100% Methanol and stored at −20°C overnight (OV). After rehydration, embryos were washed in PBBT (PBS with 2mg/mL BSA and 0.1% TritonX-100) and blocked in PBBT with 10% goat serum at RT. Subsequently, embryos were incubated OV at 4°C with primary antibodies - 1:50 mouse anti-ZO1 antibody (33-9100, *Thermo Fisher Scientific*) and 1:200 mouse anti-gamma tubulin antibody (T6557, *Millipore Sigma*). After, embryos were washed with PBBT, followed by blocking solution, and incubated overnight at 4°C with secondary antibody - 1:300 anti-mouse Alexa Fluor 546 IgG (A-11030, *Thermo Fisher Scientific*). Embryos were finally washed with PBBT and stored in PBS at 4°C. For imaging, single embryos were flat mounted onto the dish and imaged in a TCS SP8 confocal microscope (*Leica Microsystems*).

### Quantifications and statistical analysis

To statistically test the mirror-symmetry in the KV, we used a permutation test (also called randomization test) (Hesterberg T, 2005). We compute the statistic |φ_avg_left_ − φ_avg_right mirror_|, where φ_avg_left_ is the φ angle of the average resultant vector of left cilia and φ_avg_right mirror_ is the φ angle of the resultant vector of right cilia after LR mirror symmetry. This statistic is based on the definition of chirality as we test if the left cilia orientation superimposes with the mirror image of right cilia. Left and right labels of cilia are then randomly permutated 300,000 times to construct the sampling distribution of possible |φ_avg_left_ − φ_avg_right mirror_| values. The p-value is finally estimated as the proportion of permutations resulting in values greater than or equal to the experimental one. We define the structure of the KV as chiral (or asymmetric) when the p-value is lower than 0.05 and the null hypothesis (φ_avg_left_ = φ_avg_right mirror_) can be rejected. The same test is used to investigate θ mirror-symmetry. The p-values of the effective angular velocity vector 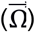 of all conditions against the WT were calculated using Welch’s test on the dorsal component Ω_D_.

### Fluid dynamic simulations

We characterized the circulatory flow in the KV by calculating the effective angular velocity vector 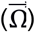 as described in (Ferreira et al., 2017). We described each cilium in a KV as a chain of beads moving along a tilted cone with the orientation obtained from 3D-CiliaMap and calculated the flow using Green’s function for the Stokes equation in the presence of a spherical no-slip boundary. The effective 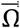 is defined as the angular velocity of a rotating rigid sphere with the same angular momentum as the time-averaged flow in the KV.

### Analysis of cilia basal body positions

Embryos at 8SS were fixed by MEMFA and labeled with anti-ZO1 and anti-gamma tubulin antibodies, followed by Alexa Fluor 546 IgG labeling (detailed protocol in ‘Immunohistochemistry’ method section). Samples were imaged in a TCS SP8 confocal microscope (*Leica Microsystems*). Cilia basal bodies were segmented in 3D from fluorescence images using Imaris (Bitplane Inc.). A local reference frame at the origin of each basal body was defined to identify the local tangent plane to the vesicle in 3D. Using custom-made scripts in Matlab (The MathWorks Inc.), the fluorescence intensity of pixels up to 2μm away from it was orthogonally projected on this plane (Sup. Figure 3B) to manually draw the cell contour and extract the antero-posterior and left-right extension of the cell. The relative position of the basal body relatively to them has then been calculated for every cilium as shown in Sup. Figure 3B-C.

## Supplementary figures

**Sup. Figure 1:**
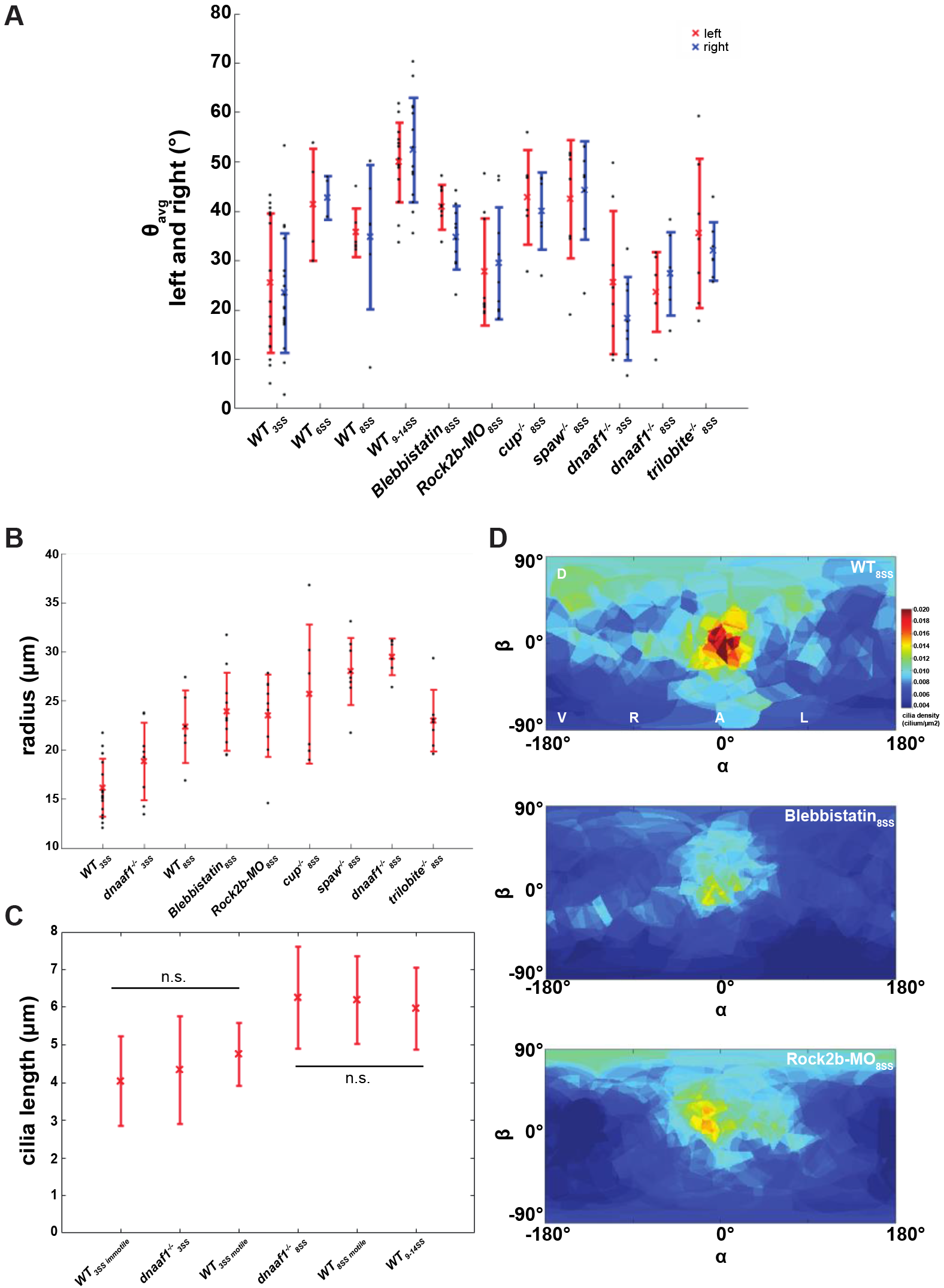
Distribution of θ angles on the right and on the left sides of the KV and additional information about the KV size, cilia length and cell density in different experimental conditions. **(A)** Dot plot displaying the mean values of θ_avg_ on the left (in red) and on the right (in blue) sides of the KV for all conditions. Permutation tests show there are no signs of asymmetry relating to θ_avg_ in all conditions studied (p > 0.05). **(B)** Average KV radius for WT and other conditions analyzed, at 3 and 8SS. The average KV radius shows no significant difference between conditions from the same developmental stage. **(C)** Average cilia length of WT immotile and motile cilia and *dnaaf1*^−/−^ immotile cilia, at 3- and 8SS. There is no significant difference between the average cilia length of WT and *dnaaf1*^−/−^ KV cilia. **(D)** Averaged cilia density obtained from KVs of WT, blebbistatin, and rock2b-MO represented on a 2D flat map revealing a disruption of the steep density gradient along the anteroposterior (AP) axis observed in WT (enrichment at the anterior pole - in red).

**Sup. Figure 2:**
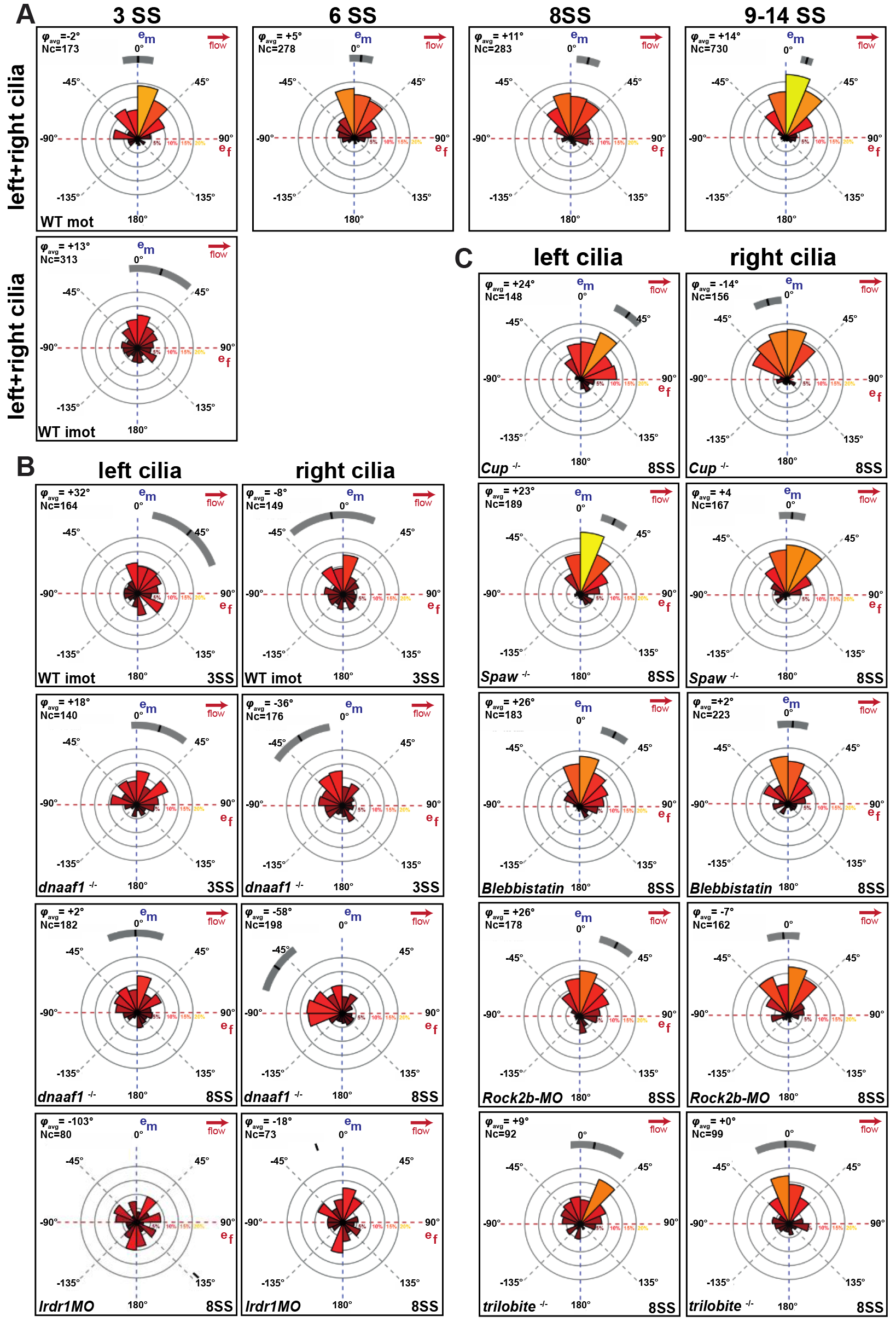
Additional information about φ angle distributions displayed in rosette plots. Rosette plots showing the φ angle distribution and φ_avg_ values of cilia in both KV hemispheres (left+right cilia) for WT cilia at different stages **(A)**, and left and right separately for the immotile cilia population in both WT, *dnaaf1*^−/−^ and *lrdr1*-MO **(B)** and for the *cup*^−/−^, *spaw*^−/−^, blebbistatin, *rock2b*-MO and *trilobite*^−/−^ conditions (C). All conditions presented asymmetric KVs at 8SS, with the exception of *trilobite*^−/−^, which is mirror-symmetric like KVs analyzed at 3SS and 6SS. Statistics provided in Table 1. Nc = number of cilia

**Sup. Figure 3:**
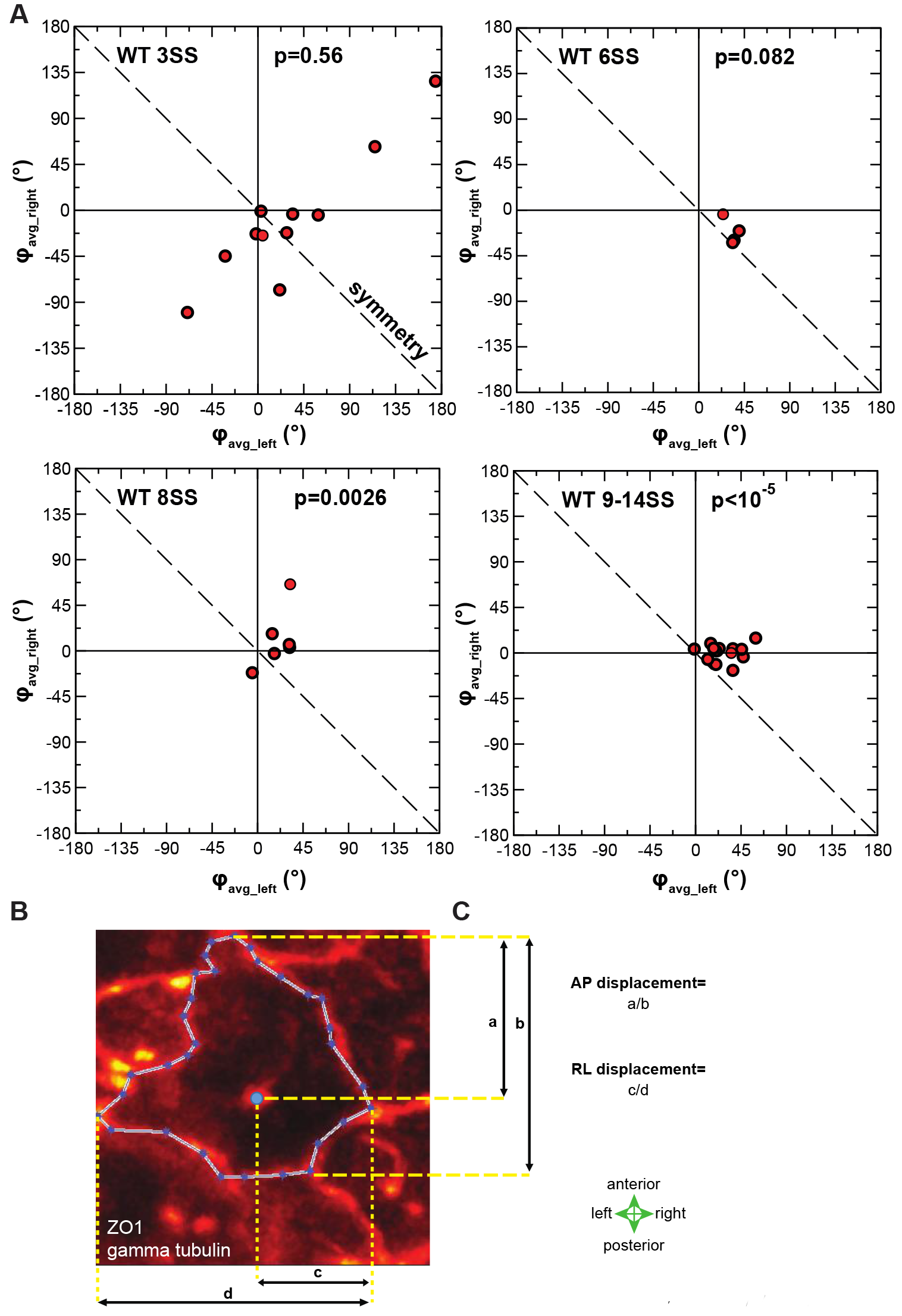
Asymmetry of cilia orientations in individual WT embryos at different stages and basal body localization quantification. **(A)** Plot showing the average orientation (φ_avg_) in the right- vs. the left half of the KV at a given stage. In a symmetric KV, the points would lie on the dashed line, indicating φ_avg_right_=−φ_avg_left_. Although the variability among embryos remains large, all 14 KVs at the later stage display an asymmetry with the same sign. The p-values (obtained from a permutation test) indicate the likelihood that an intrinsically symmetric KV (null hypothesis) would randomly show the same or larger degree of asymmetry. **(B-C)** Quantification of basal body position relative to the KV cell extent. **(B)** Orthogonal projection of the fluorescence intensity on the plane tangential to the KV surface. Basal body position (large blue dot) and cell contour (small blue dots connected with white lines) are used to measure the anterior-to-posterior and right-to left cell extents (b and d values, respectively) and the anterior-to-basal body and right-to-basal body distances (a and c values, respectively). **(C)** The anterior-to-posterior (AP) and right-to-left (RL) basal body relative positions are then estimated as a/b and c/d, respectively.

**Supplemental Figure 4:**
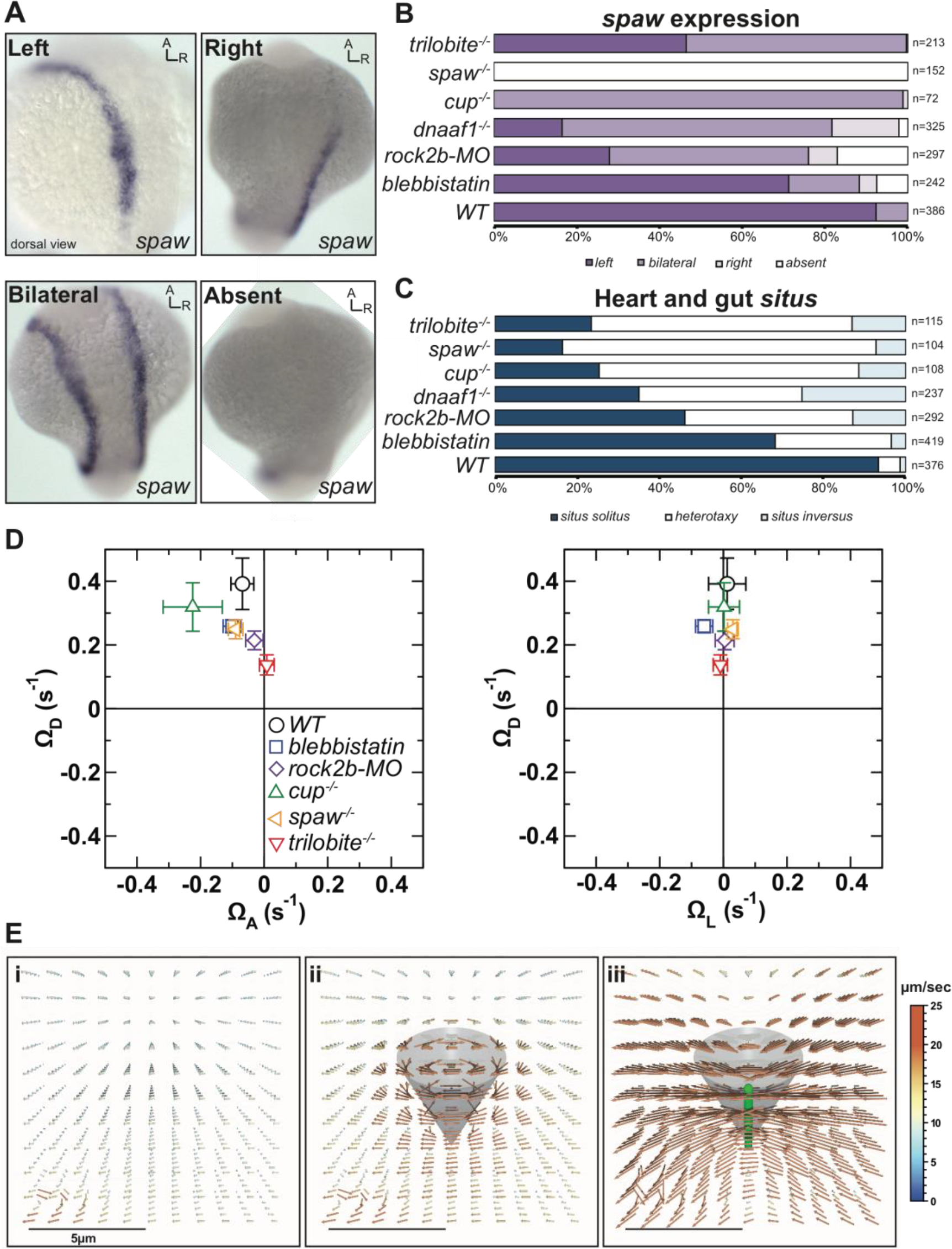
Quantification of the *spaw* expression patterns in the LPM and heart and gut *situs* in all conditions studied and flow profiles around simulated beating cilia. **(A)** *spaw* expression patterns in the LPM can be divided in left, bilateral, right or absent. **(B-C)** Percentages of *spaw* expression patterns in the LPM **(B)** and the *situs* phenotypes **(C)** respectively. The number of embryos analyzed is displayed next to each bar. *Situs* phenotypes are classified according to the clinical terminology in *situs solitus, heterotaxy* and *situs inversus*. **(D)** Effective angular velocity (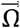 of the flow in the KV, determined computationally from cilia orientations and motility. The main graph shows the right view of the 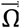 (dorsal vs. left anterior component), the insets show the anterior view (dorsal vs. left component). In contrast with the other conditions, *trilobite*^−/−^ embryos have a significantly weaker flow (p=0.024) when compared to WT at 8SS. **(E)** Flow around a cilium located at the right pole of the vesicle, determined in a simulation with a randomly generated (“synthetic”) cilia distribution, see (Ferreira et al., 2017) viewed from the center of the KV: i) time-averaged flow if the central cilium is removed and all others preserved, such that only the global flow remains, ii) time-averaged flow, iii) instantaneous flow. The results show that the global circulatory flow is much weaker than the local flows around a moving cilium.

**Movie 1: Cilia motility is impaired in the *dnaaf1*^−/−^ embryos.** The movie shows the 3D live imaging of two KVs from the *Tg*(*dnaaf1*^*tm317b*^; *actb2*:Mmu*.Arl13b-GFP*) (Sullivan-Brown et al., 2008). Each embryo was soaked for 60 minutes in Bodipy TR (Molecular Probe) and imaged using 2PEF microscopy at 930 nm wavelengths, as described in (Ferreira et al., 2017). A full z-stack of both KVs can be seen. On the left, a KV from a sibling *dnaaf1*^+/+^ with motile cilia (fan cones) is shown. On the right, a KV from a *dnaaf1*^−/−^ embryo, with 100% of immotile cilia (bright straight lines) is shown.

## Acknowledgements

We thank C. Norden, M. Blum, M. Fürthauer and the Vermot lab for discussion and thoughtful comments on the manuscript, in particular R. Chow for her help with editing. We also thank the Heisenberg lab for sharing the *trilobite* line. This work was supported by FRM (DEQ20140329553), the ANR (ANR-15-CE13-0015-01, ANR-12-ISV2-0001-01), the EMBO Young Investigator Program, the European Community, ERC CoG N°682938 Evalve and by the grant ANR-10-LABX-0030-INRT, a French State fund managed by the Agence Nationale de la Recherche under the frame program Investissements d’Avenir labeled ANR-10-IDEX-0002-02. R.R.F. was supported by the IGBMC International PhD program (LABEX). A.V. acknowledges support from the Slovenian Research Agency (grant P1-0099).

